# A size principle for bistability in mouse spinal motoneurons

**DOI:** 10.1101/2023.09.29.559784

**Authors:** Ronald M Harris-Warrick, Emilie Pecchi, Benoît Drouillas, Frédéric Brocard, Rémi Bos

**Author notes:** These authors contributed equally (Frédéric Brocard, Rémi Bos).

## Abstract

Bistability in spinal motoneurons supports tonic spike activity in the absence of excitatory drive. Earlier work in adult preparations suggested that smaller motoneurons innervating slow antigravity muscle fibers are more likely to generate bistability for postural maintenance. However, whether large motoneurons innervating fast-fatigable muscle fibers display bistability related to postural tone is still controversial. To address this, we examined the relationship between soma size and bistability in lumbar ventrolateral α-motoneurons of ChAT-GFP and Hb9-GFP mice across different developmental stages: neonatal (P2-P7), young (P7-P14) and mature (P21-P25). We found that as neuron size increases, the prevalence of bistability rises. Smaller α-motoneurons lack bistability, while larger fast α-motoneurons (MMP-9^+^/Hb9^+^) with a soma area ≥ 400µm^2^ exhibit significantly higher bistability. Ionic currents associated with bistability, including the persistent Nav1.6 current, thermosensitive Trpm5 Ca^2+^-activated Na^+^ current and the slowly inactivating Kv1.2 current, also scale with cell size. Serotonin evokes full bistability in large motoneurons with partial bistable properties, but not in small motoneurons. Our study provides important insights into the neural mechanisms underlying bistability and how motoneuron size dictates this process.

**New and Noteworthy:** Bistability is not a common feature of all mouse spinal motoneurons. It is absent in small, slow motoneurons but present in most large, fast motoneurons. This difference results from differential expression of ionic currents that enable bistability, which are highly expressed in large motoneurons but small or absent in small motoneurons. These results support a possible role for fast motoneurons in maintenance of tonic posture in addition to their known roles in fast movements.

## Introduction

Spinal motoneurons not only transmit central commands for movement to muscles but also shape motor output and muscle contraction through nonlinear firing properties (1). One such property is bistability, where the motoneuron can switch between silent and active states, depending on synaptic input or current injection. Originally detected as plateau potentials in invertebrate neurons (2–9), bistability was soon found in vertebrates (10), specifically in spinal motoneurons (11–16). While often induced by neuromodulators such as serotonin (13, 15–17), bistability can also arise when motoneurons are depolarized to near threshold and recorded under natural ionic and temperature conditions (18, 19).

The active state in bistable motoneurons is mainly mediated by slow ionic currents including persistent Cav1.3 (13–16) and Nav1.6 current (20, 21) Trpm5-mediated Ca^2^-activated Na^+^ current (18, 19, 22). Drawing from a series of our previous investigations, we can summarize the process as follows: The initial depolarization, caused by the slow inactivation of Kv1.2 potassium channels (23), activates the persistent Nav1.6 current leading to spiking activity (Drouillas et al., submitted). This then prompts Ca^2+^ entry through the recruitment of Cav1.3 channels, initiating a Ca^2+^-induced Ca^2+^-released process (18). This process ultimately activates thermosensitive Trpm5 channels, which are the primary source of the plateau depolarization to sustain repetitive firing (19). Other currents such as the HCN-type hyperpolarization-activated inward current, I_h_ (24, 25) and reduction of calcium-activated outward currents (17, 24) are also involved in different neurons.

Despite extensive studies, it remains unclear whether all motoneurons exhibit bistability. In decerebrate cats, small motoneurons characterized by slow conduction velocities and low activation thresholds, appear to have more pronounced bistability than large ones (15, 16). This observation suggests that motoneurons innervating slow (S) or fast fatigue-resistant (FR) muscle fibers show higher bistability than those linked to fast fatigable (FF) muscle fibers. Notably, S motoneurons, primarily associated with postural adjustments and slow movements, are believed to use their bistability for efficient postural maintenance, ensuring minimal energy expenditure (15, 16, 26).

To further investigate the size distribution of bistability, we conducted a study on genetically labeled α-motoneurons in young mice (postnatal day (P) 1-25). Unlike the cat studies(15, 16), we found that the largest α-motoneurons showed stronger bistable firing properties, while the smallest neurons were rarely bistable. There was a strong correlation between bistability and the amplitudes of the ionic currents known to support it. Motoneurons with intermediate sizes often showed partial bistability, which could be converted to full bistability by serotonin. These unexpected results lead to a reappraisal of the role of larger motoneurons in bistability and postural control.

## Materials and methods

### Animals and preparation

Hb9-GFP mice were kindly provided by B. Pettmann and Jackson Laboratories (strain 005029). ChAT-Cre mice were obtained from Jackson Laboratories (strain 007902) and crossed to Rosa-26-floxed GFP mice. Cornell: All animal protocols were approved by the Cornell Institutional Animal Use and Care Committee and were in accordance with NIH guidelines. Marseille: All animal care and use conformed to French regulations (Décret 2010-118) and were approved by the local ethics committee (Comité d’Ethique en Neurosciences INT-Marseille, CE71 Nb A1301404, authorization Nb 2018110819197361). Animals were housed on a 12 hr day/night cycle with ad libitum access to water and food. The room temperature was maintained between 20 and 21°C.

Mice were cryoanaesthetized (P2-P7) or anaesthetized (P8-P25) with intraperitoneal injection of a mixture of ketamine/xylazine (100mg/kg and 10 mg/kg, respectively). They were then decapitated, eviscerated and the spinal cord removed by laminectomy, and placed in ice cold (1-2°) aCSF containing (in mM): 252 sucrose, 3 KCl, 1.25 KH_2_PO_4_, 4 MgSO_4_, 0.2 CaCl_2_, 26 NaHCO_3_, 25 D-glucose, pH 7.4, bubbled with 95% O_2_ and 5% CO_2_. The meninges were removed and the posterior cord (L3-S5) imbedded in agarose. The same solution was used for slicing. 325-350µm sections were prepared from the L4-L5 region and transferred to recording aCSF containing (in mM): 120 NaCl, 3 KCl, 1.25 NaH_2_PO_4_, 1.3 MgSO_4_, 1.2 CaCl_2_, 25 NaHCO_3_, 20 D-glucose, pH 7.4, 32-34°C. In most experiments (P1-P14), blockers of fast synaptic transmission (CNQX and D,L-AP5 or kynurenic acid, strychnine, and bicuculline) were added in the aCSF to minimize synaptic contributions to bistability. To isolate the persistent Na^+^ currents (INaP) during voltage-clamp experiments we used a modified aCSF containing (in mM): 100 NaCl, 3 KCl, 1.25 NaH_2_PO_4_, 1.3 MgSO_4_, 3.6 MgCl_2_, 1.2 CaCl_2_, 25 NaHCO_3_, 40 D-glucose, 10 TEA-Cl and 0.1 CdCl_2_.

Mature mice (P21-25) were treated as the younger mice, except that the cord was sliced in an ice cold slicing solution (1-2°) containing (in mM): 130 K-gluconate, 15 KCl, 0.05 EGTA, 20 HEPES, 25 D-glucose, 3 kynurenic acid, and pH 7.4 with NAOH (27). After a resting period of 30-60 min, slices were transferred to the recording chamber and superfused with recording aCSF at 32°C (28) without addition of fast synaptic transmission blockers.

### Electrophysiological methods

Hb9-GFP and ChAT-CFP neurons were visualized in the ventrolateral region of lamina IX in L4-L5 slices. Whole-cell patch-clamp recordings were made using electrodes (2-6MΩ) pulled from borosilicate glass capillaries (1.5 mm OD, 1.12 mm ID; World Precision Instruments). They were filled with a pipette solution containing (in mM): 140 K^+^-gluconate, 5 NaCl, 2 MgCl_2_, 10 HEPES, 0.5 EGTA, 2 ATP, 0.4 GTP, pH 7.3. Patch clamp recordings were made using a Multiclamp 700B amplifier driven by PClamp 10 software (Molecular Devices). Recordings were digitized on-line and filtered at 10 kHz (Digidata 1322A or 1440A, Molecular Devices). Pipette and neuronal capacitive currents were canceled, and, after breakthrough, series access resistance was compensated and monitored. The recording was allowed to stabilize for at least 2 min after establishing whole-cell access before recording was started.

### Drug list

All solutions were oxygenated with 95% O_2_/5% CO_2_. All salt compounds, as well as tetraethylammonium chloride (TEA; #T2265), Triphenylphosphine oxide (TPPO; #T84603), Serotonin creatinine sulfate monohydrate (5-HT; #H7752), 6-Cyano-7-nitroquinoxaline-2,3-dione (CNQX, #5.04914), D-(-)-2-Amino-5-phosphonopentanoic Acid (DL-AP5; #165304), strychnine (#S0532), bicuculline (#5.05875), Kynurenic Acid (#K3375) and Amphotericin B (#A4888) were obtained from Sigma-Aldrich. Tetrodotoxin (TTX; #1078) was obtained from Tocris Bioscience.

### Immunohistochemistry

Spinal cords from 10-12-day-old Hb9-GFP mice were removed and fixed for 5-6 h in 4% paraformaldehyde (PFA) prepared in phosphate buffer saline (PBS), then rinsed in PBS and cryoprotected overnight in 20% sucrose-PBS at 4°C. Spinal cords were frozen in OCT medium (Tissue Tek), and 30 μm cryosections were collected from the L4-L5 segments. After washing in PBS 3×5 min, the slides were incubated for 1 h in a blocking solution (BSA 1%, Normal Donkey Serum 3% in PBS) with 0.2% triton X-100 and for 12 h at 4 °C in a humidified chamber with the primary antibody: mouse-anti-NeuN (Neuronal Nuclei, Sigma-Aldrich MAB377) or goat-anti-MMP-9 (Matrix metallopeptidase 9, Sigma-Aldrich M9570). Both antibodies were diluted in the blocking solution with 0.2% Triton X-100 (1:1000 and 1:500 for anti-NeuN and anti-MMP-9, respectively). Slides were washed 3×5 min in PBS and incubated for 2 h with an Alexa Fluor® Plus 555-conjugated secondary antibody (Invitrogen A32816) diluted in the blocking solution. After 3 washes of 5 min in PBS, they were mounted with a gelatinous aqueous medium. Images were acquired using a confocal microscope (LSM700, Zeiss) equipped with a 40x oil objective and processed with the Zen software (Zeiss).

### Analysis and statistics

Electrophysiological data were analyzed with Clampfit 10 software (Molecular Devices). Only cells with a stable membrane potential below -60 mV, stable access resistance, and action potential amplitude larger than 60 mV were analyzed. Reported membrane potentials were not corrected for liquid junction potentials. Statistical analysis was carried out mainly using GraphPad Prism and Matlab (MathWorks) software. Fisher’s exact test, two-tailed paired and unpaired t-tests were used as needed; p values <0.05 were considered significant. Each statistical test is indicated in the figure legends. In the figures, data are presented as mean ± SEM for the histograms. Median and quartiles are represented in each violin plot.

## Results

### Large ventrolateral spinal neurons are fast α-motoneurons

Motoneurons in the ventrolateral spinal cord were initially identified by their expression of choline acetyltransferase (ChAT) in ChAT-GFP mice. We found a number of labeled neurons of many sizes (data not shown); these include both α-motoneurons driving extrafusal muscle fibers and smaller γ-motoneurons innervating intrafusal muscles that regulate muscle spindles’ responsiveness to stretch. These motoneuron subtypes can be distinguished by differential expression of a number of proteins. While the transcription factor Hb9 is a marker for both α- and γ-motoneuron in Hb9-nls-LacZ animals (29), only α-motoneurons express strong GFP labeling in the Hb9-GFP mice (29). We thus used Hb9-GFP mice to identify α-motoneurons in the L4-L5 ventrolateral spinal cord, preferentially linked to extensor muscles, from postnatal day 1 (P1) to P25.

The transcription factor NeuN is commonly used to distinguish neurons from glial cells. In the spinal cord, α-motoneurons strongly express NeuN, whereas γ-motoneurons either lack expression or exhibit weak expression in 2/3 and 1/3 of cases, respectively (30). We examined co-expression of Hb9-GFP and NeuN immunolabeling and, regardless of age, we found that a significant majority of ventrolateral Hb9-GFP-labeled neurons exhibit strong NeuN labeling (98%, n = 144 cells), indicating they are likely α-motoneurons (Fig. 1A, B1). We observed a preponderance of small labeled α-motoneurons during P4-6 (n = 571 neurons; Fig. 1B2) and P8-10 (n = 425 neurons; Fig. 1B2), with 80.5% and 79.6% respectively having a maximal cross sectional area of 400 µm^2^ or less. The percentage of large α-motoneurons increases with age (29, 30), and by P12, only 50.4% of them remained under 400µm^2^ (n = 388 neurons; Fig. 1B2).

**Figure 1:**
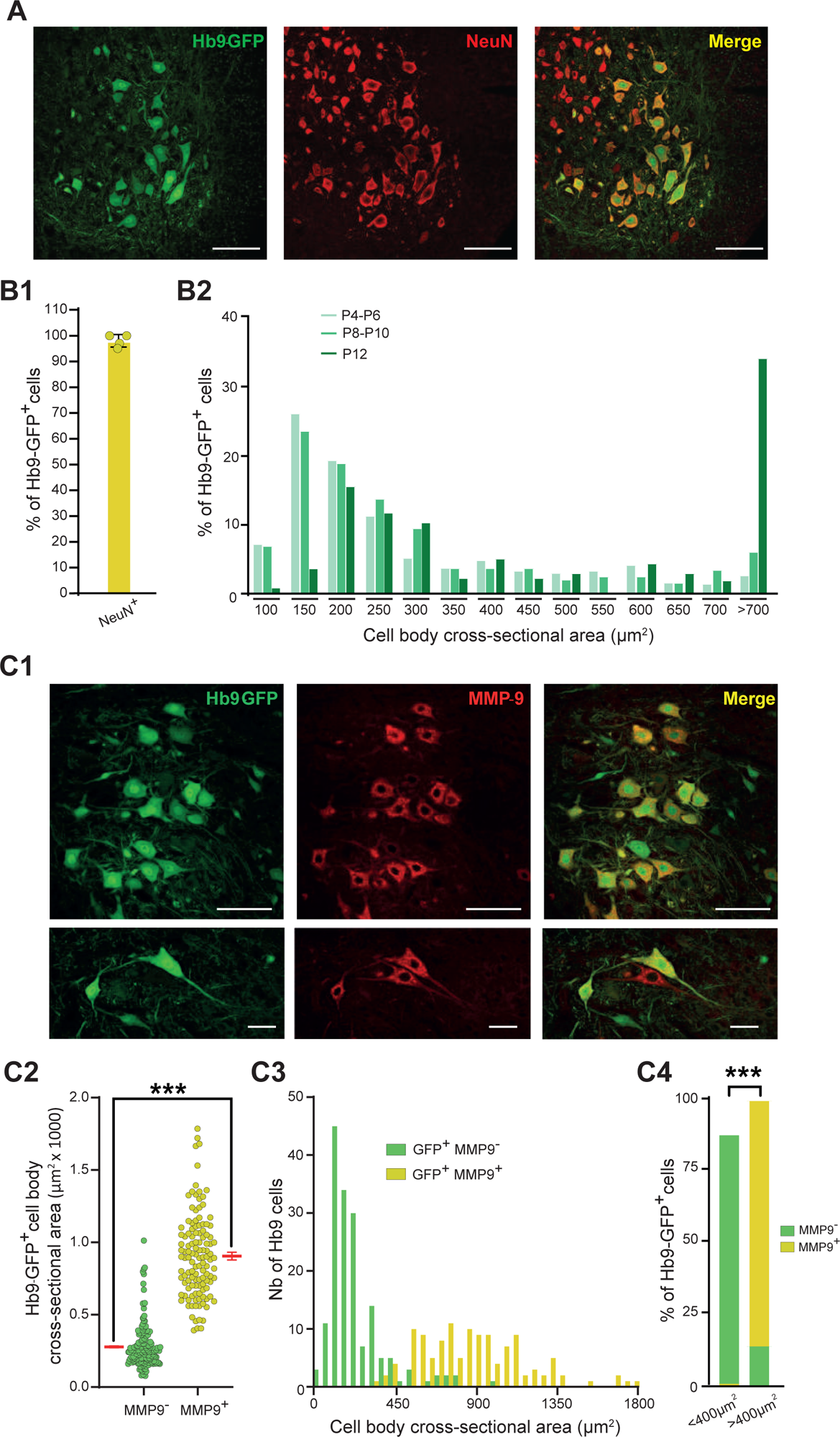
Identification and size distribution of α-motoneurons in the ventrolateral spinal cord at L4-L5. **A:** Ventrolateral cord at P12 showing Hb9-GFP (green), NeuN (red) and double labeling (yellow). Scale bar, 100µm. **B1**: The vast majority (∼98%) of Hb9-GFP-labeled neurons co-express NeuN. **B2**: Cross-sectional area distribution of Hb9-GFP neurons at P4-P6 (light green), P6-P10 (medium green) and P12 (dark green). **C1**: Ventrolateral cord at P12 showing Hb9-GFP (green), MMP-9 (red) and dual labeling (yellow). Scale bars, 100µm and 50µm for top and bottom images, respectively. **C2**: Size distribution of Hb9-GFP labeled neurons expressing or not expressing MMP-9. **C3**: Histogram of size distribution of GFP^+^-MMP-9^-^ (green) or HB9^+^-MMP-9^+^ (yellow) neurons. **C4**: Double-labeled Hb9-GFP/MMP-9 neurons are predominantly larger than 400 µm^2^ (yellow, right) while most Hb9-GFP^+^ neurons smaller than 400 µm^2^ do not co-express MMP-9 (green, left). *** p<0.001 (two-tailed Mann-Whitney test for **C2**; Fisher’s exact test for **C4**). Mean ± SEM.

α-motoneurons can be subdivided functionally into three classes based on size and the muscle type they innervate: large fast-fatigable (FF), medium fast fatigue-resistant (FR), and small slow (S) [reviewed by (31)]. Only FF motoneurons express the marker matrix metalloproteinase-9 (MMP-9) (32, 33). Our observations confirmed that larger motoneurons co-expressed Hb9-GFP and MMP-9, while the smaller ones did not (Fig. 1 C1-C4). The mean cross-sectional area of MMP-9-negative neurons was 280 ± 12 µm^2^, while that of MMP-9-positive neurons was 907 ± 26 µm^2^ (P<0.001, n = 288; Fig. 1C2). As suggested by earlier work (29), most of the motoneurons under 400µm^2^ are HB-9^+^/MMP-9^-^ (86.75%) while those over 400µm^2^ are HB-9^+^/MMP-9^+^ (99.2%; Fig. 1C3, C4). Single cell RT-PCR measurements further validated this distinction, where the larger neurons synthesized RNA for MMP-9 (n= 6 cells) while the smaller neurons or glial cells did not (data not shown), indicating larger neurons as FF motoneurons and smaller ones as S or FR motoneurons. Markers to selectively identify FR motoneurons in slice recordings performed *ex vivo* are currently unavailable.

### Features of bistable motoneurons

We previously showed that under experimental conditions mimicking natural *in vivo* conditions, many motoneurons exhibit bistability in the absence of added neuromodulators. This is achieved by maintaining *in vivo* calcium concentrations (1.2 mM) (34) and keeping the preparation temperature above 30°C (18, 19, 35, 36). Four distinct features characterize bistable motoneurons (Fig. 2): 1) self-sustained firing triggered by a brief (2 sec) excitation when the motoneuron is pre-depolarized near the spike threshold (Fig. 2A1); 2) a slow afterdepolarization (sADP) following the current step if the motoneuron is not pre-depolarized sufficiently to trigger the self-sustained firing (Fig. 2A1); 3) negative hysteresis during slow triangular current ramp injections, where spiking stops at lower currents than where it began (Fig. 2A2); 4) a slowly depolarizing potential causing delayed spiking acceleration in response to a near-threshold depolarizing step (Fig. 2A3). We assigned each features one point, with fully bistable motoneurons scoring 4 points. To be considered bistable, the motoneuron must score at least 3 points, including the presence of a self-sustained firing and sADP, and either negative hysteresis or delayed firing.

**Figure 2:**
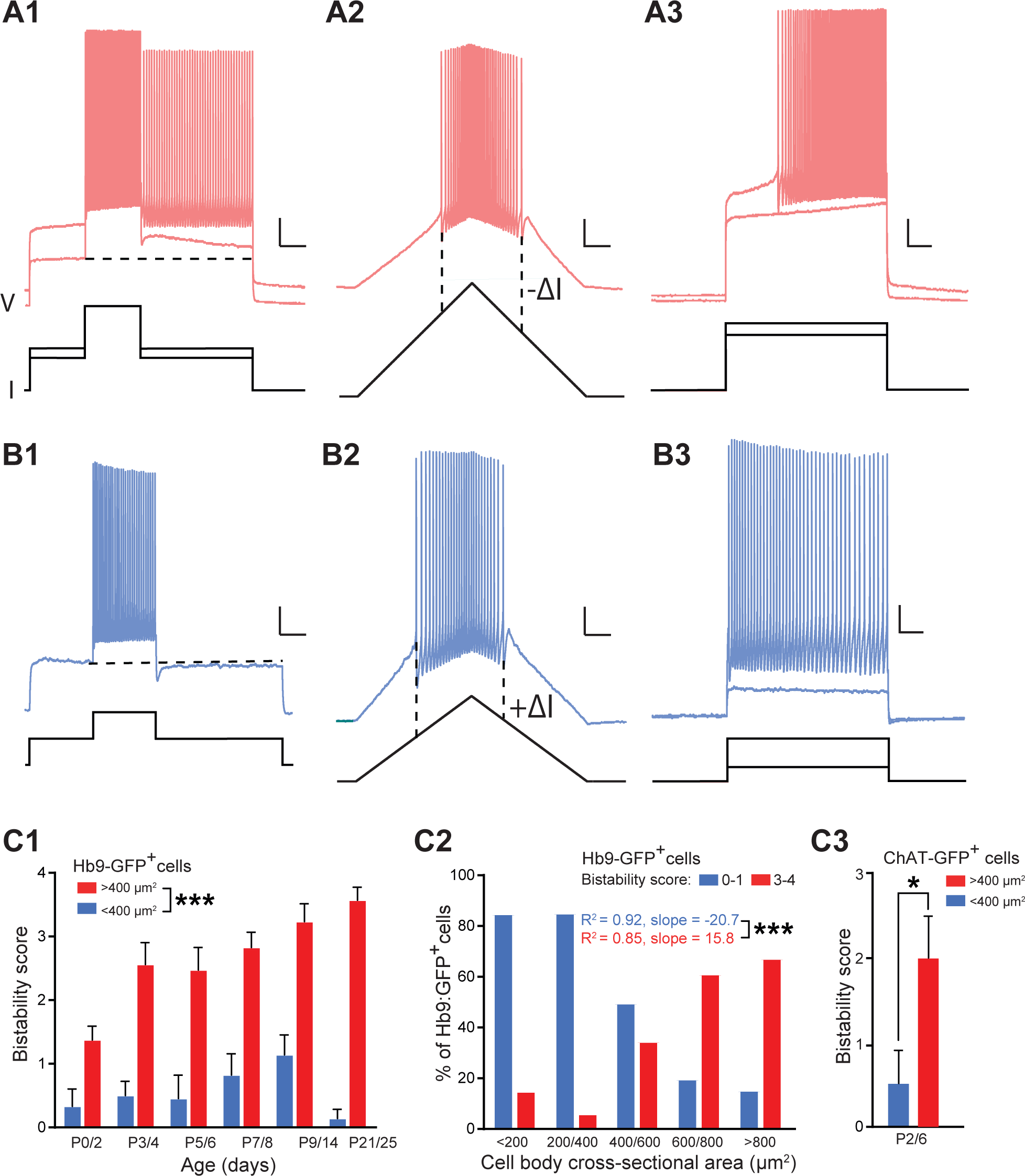
Bistability score in spinal motoneurons varies by age and size. **A**: Fully bistable neuron (bistability score 4). **A1**: Baseline depolarization towards spike threshold: a 2-sec suprathreshold stimulation evokes a prolonged afterdepolarization (ADP: 1 point). Slightly higher baseline depolarization followed by a 2 sec stimulation leads to prolonged firing which is only terminated by hyperpolarizing current step (1 point); trace has been offset down to be more easily seen. **A2**: Negative hysteresis during ramp stimulation. The current threshold for onset of spiking is higher than the threshold for offset of spiking (-ΔI; 1 point). **A3**: A small subthreshold current step leads to a slow depolarization (trace has been offset down to be more easily seen). A slightly higher current step leads to a larger depolarization leading to delayed spike onset and acceleration of spike frequency during the step (1 point). **B**: Completely non-bistable neuron (bistability score 0). **B1**: Current steps to near threshold lead to decelerating spike frequency during a 2 sec suprathreshold pulse, and an afterhyperpolarization at the end of the step (0 point). **B2**: Positive hysteresis during ramp stimulation. The current threshold for onset of spiking is lower than the threshold for offset of spiking (+ΔI; 0 point). **B3**: A small subthreshold current step does not evoke a slow depolarization. A slightly higher current step evokes immediate onset firing with decelerating spike frequency during the step (0 point). Size markers: 1 sec, 20 mV. **C1**: Bistability score as a function of postnatal age for smaller (<400µm^2^, blue) and larger (>400µm^2^, red) Hb9-GFP^+^ neurons. **C2**: Distribution of bistability as a function of neuronal cross-sectional area. Smaller neurons predominantly have bistability scores of 0-1 (blue) while larger neurons predominantly have bistability scores of 3-4 (red). **C3**: Post-natal (P2/6) motoneurons identified by expression of ChAT-GFP show similar bistability scores that rise with cross-sectional area (blue for <400µm^2^, and red for >400µm^2^). Size markers: 1 sec, 20 mV. * p<0.05; *** p<0.001 (two-tailed Mann-Whitney test for **C1** and **C3;** slope comparison of simple linear regressions for **C2**). Mean ± SEM.

Many motoneurons did not express any of the bistability criteria, scoring 0. They typically showed a decelerating firing rate during the 2 sec depolarizing pulse, leading to a post-step afterhyperpolarization (Fig. 2B1). Their activity during ramp current injections showed positive hysteresis, where spiking ceases at higher current values than where it started (Fig. 2B2). They also began spiking immediately during a suprathreshold long step, with a continuous firing deceleration (Fig. 2B3). Some motoneurons, despite having one scoring feature, lacked self-sustained firing and were also considered as non-bistable. A significant number of motoneurons also displayed intermediate characteristics, scoring 2. The vast majority of them (90 %) were not bistable, lacking self-sustained spiking, but meeting two other criteria. This shows that bistability is not an all-or none property, but can manifest with somewhat different properties.

### Effects of size and age on bistability

As previously described, bistability in motoneurons can emerge early in development (18). Fig. 2C1 shows the average bistability scores of Hb9-GFP^+^ α-motoneurons during the initial postnatal month, comparing small neurons (<400µm^2^) to large neurons (>400µm^2^). Small motoneurons consistently showed minimal bistability scores across all developmental ages (Fig. 2C1), rising up to a mean of 1.14 ± 0.31 by P9-14, then nearly vanishing by early adulthood (P21-25). Only 9 out 81 small cells (<400 µm^2^) showed bistability (scores 3-4) at any age, with 55% appearing after P9. In contrast, large motoneurons (>400 µm^2^) exhibited increasing bistability, reaching near maximal levels around P9 with a mean score of 3.24 ± 0.28. Remarkably, 83% (n = 24) of these large motoneurons exhibited high bistability scores (3–4) from P9, with all displaying bistability by P21-25 (n = 7). Note that the bistability scores of large motoneurons are significantly higher compared to small ones across all ages (P<0.001, n=199).

These data underscore the pivotal role of motoneuron soma size in shaping the emergence of bistability in young mice. Fig. 2C2 shows the bistability score distribution by size across all ages. As described above, the large majority of small neurons (<400 µm^2^) were not bistable, averaging a score of 0.67 ± 0.13 (n = 81); only 11 % achieved scores of 3-4. In contrast, larger neurons (>400 µm^2^) consistently had high bistability scores (mean 2.54±0.13, n = 118; P<0.001), with the largest (>800µm^2^) being the most bistable (mean 3.06 ± 0.17, n = 44) with 75% scoring 3-4. Intermediate size neurons (400-800 µm^2^) show largely a mixture of bistable and intermediate bistability scores (mean 2.32 ± 0.17; n = 78). A clear gradient emerges: as neuron size increases, the proportion of bistable neurons rises, while the proportion of non-bistable cells decreases, especially above 400 µm^2^. The intermediate scores of 2 (not shown) fluctuate with size, perhaps reflecting maturational intermediates over time.

Similar results collected from ChAT-GFP mice reinforce the size-dependent nature of bistable properties (Fig 2C3). Before P7, small motoneurons (<400 µm^2^) showed no bistability, with an average score of 0.56 ± 0.38, n = 9. In contrast, larger neurons (>400µm^2^) showed higher bistability (score 2.1 ± 0.42, n = 16; p = 0.02). As a control measure, we used the amphotericin B perforated patch method (28) to record bistability from HB9-GFP neurons, avoiding the dialysis of neuronal contents. This method reaffirmed the trend of increasing bistability with α-motoneuron size (n = 26; data not shown).

### Currents associated with bistability

Several ionic currents have been demonstrated to support the active state during bistability. These include a persistent inward current (PIC) that may comprise both sodium and calcium components (16, 21, 37, 38), a thermosensitive Trpm5 calcium-activated inward current (19), and a slow inactivation of the Kv1.2 potassium current (23).

We measured the PIC by delivering a slow ramp depolarization in voltage clamp (Fig. 3A). The PIC amplitude was measured at the inward peak from the extrapolated passive component at the same voltage (See Fig. 3A). The PIC amplitude was very small or absent in small Hb9-GFP^+^ motoneurons (<400µm^2^: 33 ± 14 pA, n = 17; Fig. 3B), but much larger in large motoneurons (165 ± 35 pA, n = 24; p = 0.014), with its amplitude correlating positively with cell size without any change in the activation threshold (Fig. 3C). Moreover, the amplitude of the PIC was also found to be proportional to the bistability score (Fig. 3D). Specifically, non-bistable neurons (scores 0-1) had small to absent PICs (mean 11.7 ± 5.01 pA; n = 15, 9 with no PIC), whereas bistable neurons (scores 3-4), especially those that are fully bistable (score 4), exhibited a dramatic rise in the PIC amplitude (mean 198 ± 34 pA, n = 22, P < 0.001, 0 with no PIC). This increase in PIC with bistability was not merely a reflection of the greater surface area of the bigger neurons that express more channels, as the PIC density (corrected for cell area) was much smaller in non-bistable neurons (scores 0-1; 0.035 ± 0.07 pA/µm^2^, n=19) compared to bistable neurons (scores 3-4; 0.289 ± 0.22 pA/µm^2^, n = 21; p< 4×10^-5^). These findings highlight the strong association between PICs and bistability in large α-motoneurons and the absence or low amplitude of PICs in smaller non-bistable neurons.

**Figure 3:**
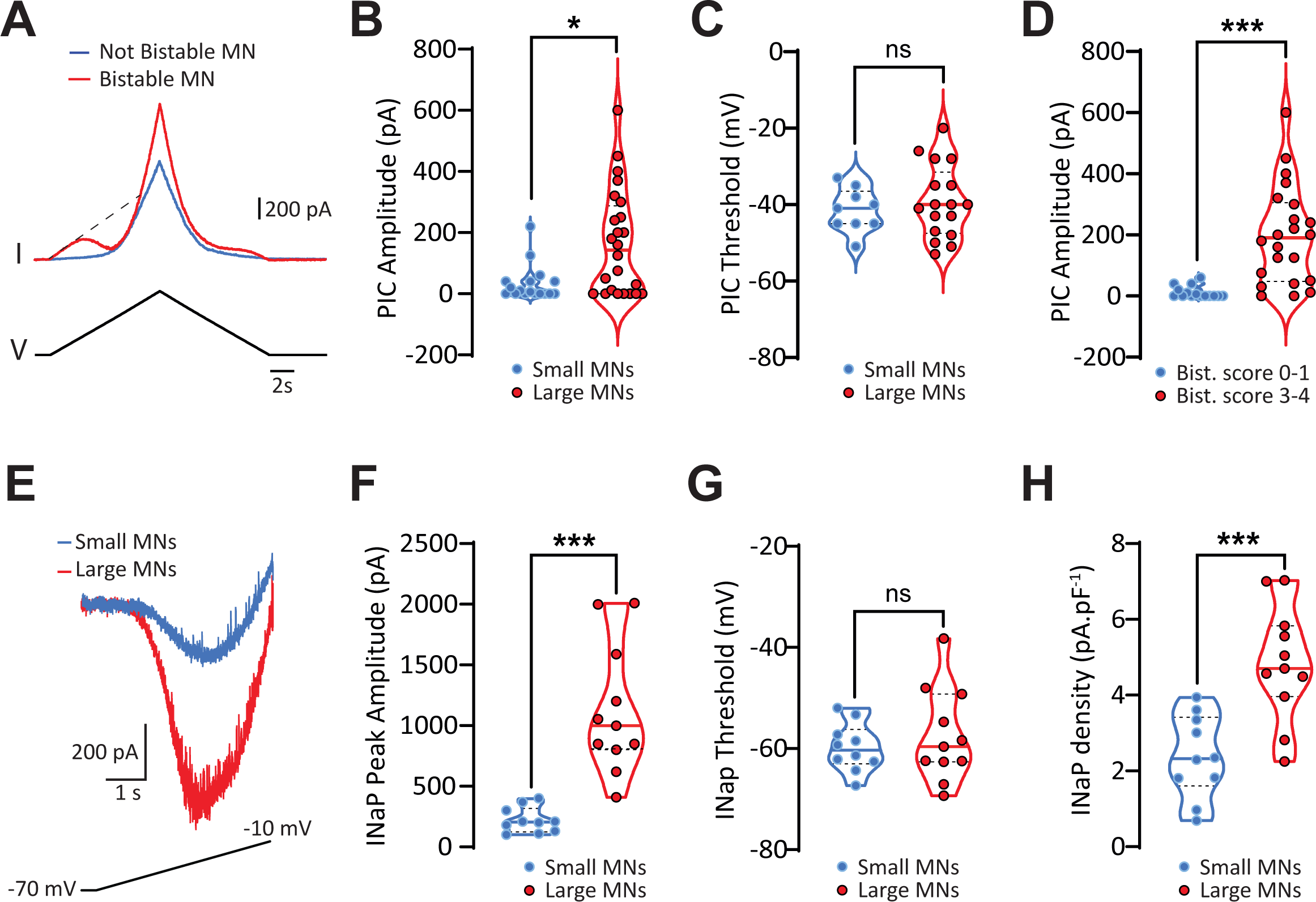
Persistent inward current (PIC), in particular its sodium component, is larger in bistable neurons. **A**: Voltage clamp measure of PIC activation during slow voltage ramp in bistable but not in non-bistable neuron. **B-C**: PIC amplitude (B), and threshold (C) as a function of cell cross-sectional area of Hb9-GFP^+^ neurons (blue circles, <400µm^2^, and red circles, >400µm^2^). **D**: PIC amplitude increases with increasing bistability (low and high bistability scores in blue and red, respectively). **E**: Superimposed leak-subtracted sodium persistent current (INaP) recorded from Hb9-GFP^+^ motoneurons in the presence of 10 mM TEA and 100µM CdCl_2_ (area <400µm^2^ or >400µm^2^ for the blue and red trace, respectively) in L4-L5 regions at P9 in response to a slow ramp depolarization. **F-H**: INaP peak amplitude (F), threshold (G) and density (H) as a function of cell cross-sectional area of Hb9-GFP^+^ neurons (blue circles, <400µm^2^, and red circles, >400µm^2^). ns, no significance; *p<0,05; ***p<0,001 (two-tailed Mann-Whitney test for **B-D** and **F-H**). Median (solid line) and quartiles (dashed lines) are represented in each violin plot.

To further understand the contribution of the sodium component of PICs in bistability, we added 10 mM TEA-Cl and 100 µM CdCl_2_ to the recording ACSF (39). Large α-motoneurons displayed a more pronounced persistent sodium current (INaP; Fig. 3E). Although the motoneuron size did not influence the INaP threshold activation, both the peak amplitude and density of INaP were significantly greater in large α-motoneurons (>400 µm^2^, 1126 ± 160 pA & 4.84 ± 0.46 pA/pF, n = 11) compared to small motoneurons (<400 µm^2^, 221 ± 33 pA & 2.38 ± 0.35 pA/pF, n = 10) (Fig. 3F-H; P < 0.001). These findings highlight the significant contribution of INaP in bistability from large α-motoneurons and its diminished presence in smaller, non-bistable neurons.

A calcium-activated inward current, mediated by Trpm5 channels, has been recognized as pivotal in sustaining the plateau depolarization underlying the tonically firing active state in bistable neurons (19). To measure the effect of this current, we induced a strong 2-sec depolarization in the presence of tetrodotoxin (TTX, 1 µM) and tetraethylammonium-chloride (TEA, 10 mM) to minimize sodium and potassium currents. This depolarization was large enough to elicit a series of slow calcium-driven spikes to fully activate Trpm5. Subsequently, we measured the resulting afterdepolarization (sADP; Fig. 4A), which slowly declined as the Trpm5 current decayed (19). Notably, the sADP was partially blocked by Triphenylphosphine oxide (TPPO, 50 µM Fig. 4A), a known Trpm5 channels blocker (19, 40). The residual sADP appears to predominantly arise from channels that are not yet identified (see discussion in (19)). Thus, to accurately estimate the Trpm5 component of the sADP, we subtracted the sADP measured in TPPO from the total sADP. The resulting Trpm5 sADP was only detectable in one of 15 smaller neurons (<400 µm^2^: 0.2 ± 0.24 mV, n = 15), but was evident in most of the larger neurons (>400 µm^2^: 4.9 ± 1.83 mV, n = 9; p = 0.0017; Fig. 4A-B). Neurons with lower bistability scores (scores 0-1) lacked Trpm5 sADP (n=10), but this current was present in most bistable neurons (scores 3-4: 4.6 ± 1.68 mV, n = 10; p = 0.0108; Fig. 4C). The larger Trpm5 sADP in more bistable currents does not simply represent the larger size (and surface area) of these neurons; when corrected for area, the Trpm5 ADP density was absent in non-bistable neurons (scores 0-1, 0.0 mV/µm^2^, n = 10) but present in more bistable neurons (scores 3-4, 7.7µV/µm^2^, n = 10; p = 0.014). We isolated the Trpm5 calcium-activated inward current in response to a 2 sec depolarizing voltage step in presence of TTX (1µM) and TEA (10mM) (Fig.4D). We confirmed that the Trpm5 inward current amplitude and density are significantly bigger in large α-motoneurons (>400 µm^2^, 324 ± 51.73 pA & 2.7 ± 0.49 pA/pF, n = 8) compared to the small ones (<400 µm^2^, 26 ± 4.33 pA & 0.7 ± 0.39 pA/pF, n = 3) (Fig. 4E-F; P<0.05). Thus, the calcium-activated inward current mediated by Trpm5 channels (19) appears associated with bistability in large α-motoneurons, but is low or absent in smaller non-bistable neurons.

**Figure 4:**
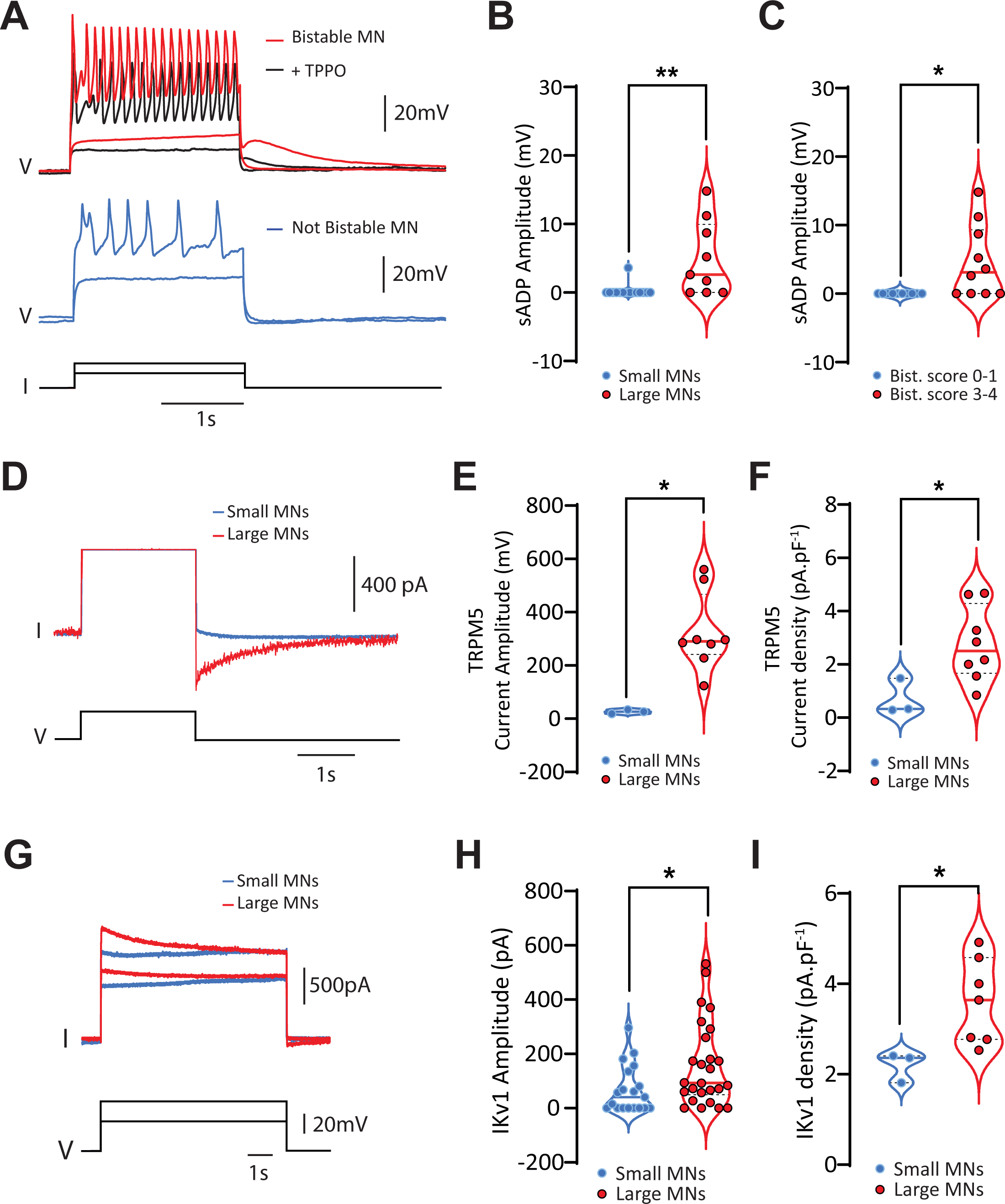
Trpm5-mediated current and slowly inactivating Kv1 current are signatures of bistability. **A**: Neurons are depolarized for 2 sec in presence of 1µM TTX and 10 mM TEA. Above a threshold, large calcium-dependent oscillations are evoked. In bistable motoneurons (red trace), these elicit a large, slow afterdepolarization which is partially blocked by 50µM TPPO (black trace). Non-bistable neurons (blue trace) do not express this slow afterdepolarization. **B**: Amplitude of the Trpm5-induced, TPPO-resistant afterdepolarization increases with cross-sectional area. **C**: Trpm5-induced, TPPO-resistant afterdepolarization increases with bistability. **D**: TPPO-sensitive Trpm5 current is isolated in response to a voltage step of 2 sec in presence of 1µM TTX and 10 mM TEA. Superimposed traces of Trpm5 isolated current from large (>400µm^2^, red) *vs* small (<400µm^2^, blue) Hb9-GFP^+^ motoneurons. **E-F**: Amplitude (E) and density (F) of the Trpm5 current increases with cross-sectional area of Hb9-GFP^+^ neurons (blue circles, <400µm^2^, and red circles, >400µm^2^). **G**: Superimposed traces of the slowly inactivating Kv1 current in bistable (red) and non-bistable (blue) neuron in response to a long depolarizing voltage step in presence of 1µM TTX and 10 mM TEA. **H-I**: Amplitude (H) and density (I) of Kv1 current increases with cross-sectional area of Hb9-GFP^+^ neurons (blue circles, <400µm^2^, and red circles, >400µm^2^). *p<0,05; **p<0,01 (two-tailed Mann-Whitney test for **B-C**, **E-F** and **H-I**). Median (solid line) and quartiles (dashed lines) are represented in each violin plot.

We further assessed the Kv1.2 potassium current, whose slow inactivation delays the initiation and acceleration of firing of bistable neurons during long current steps (23). This was measured in voltage clamp as the amplitude of the inward current over a 7 sec step, measured after inactivation of the transient potassium current, in presence of TTX (1µM) and TEA (10mM) (Fig. 4G). Again, the amplitude of the Kv1.2 current scaled with cell size. Smaller α-motoneurons (<400 µm^2^) had a lower current amplitude (52.5 ± 16.8 pA; n = 16) than the larger neurons (135.6 ± 27.7 pA, n = 23; P < 0.02; Fig. 4H). Moreover, these smaller motoneurons has a lower current density (2.19 ± 0.33 pA/pF; n = 3) compared to larger motoneurons (3.61 ± 0.35 pA/pF, n = 7; P < 0.05; Fig. 4 I).

All together, our findings highlight that INaP, the Trpm5 mediated calcium-activated inward current, and the slow inactivating Kv1 current are pivotal indicators of bistability in large α-motoneurons.

### Serotonin evokes bistability only in larger, partially bistable neurons

We found that bistability is not strictly binary, as many α-motoneurons displayed some, but not all, of the defining features described above, especially the absence of persistent firing after the brief depolarization. It has been known for many years that neuromodulators such as serotonin (5**-**HT) can evoke bistability in α-motoneurons which show only partial bistable properties (14, 17, 24, 41).

We analyzed the effects of 10 µM 5-HT on α-motoneurons (n = 52) that had bistability scores below 4 and lacked the self-sustained spiking. Fig. 5A shows an example of an α-motoneuron that, under control conditions (i.e. recording aCSF), did not show acceleration of spiking during a 2-sec strong depolarization, and only showed a modest sADP at the end of the depolarization. Seven minutes after application of 5-HT, this motoneuron demonstrated accelerating spiking during the current step, and self-sustained spiking activity after the end of the step. This activity was terminated only by a hyperpolarizing step, indicating that the neuron became fully bistable only in the presence of 5-HT. Fig. 5B shows the ability of 5-HT to evoke bistability in α-motoneurons of different sizes. Notably, small α-motoneurons (< 400µm^2^; n = 19) remained non-bistable even with 5-HT. In contrast, larger neurons exhibited the potential for a bistable transition upon 5-HT exposure, and this ability increased with cell size. Specifically, of non-bistable α-motoneurons of intermediate size (400-800µm^2^), 25% switched to full bistability with 5-HT (n = 28). Of the few larger α-motoneurons (> 800µm^2^) which were not initially bistable, 60% became fully bistable with 5-HT (n = 5; P < 0.001). It’s noteworthy that the majority of neurons transitioning to bistability with 5-HT already expressed an sADP before 5-HT addition, as illustrated in Fig. 5A. Thus, 5-HT is able to evoke full bistability in a subset of neurons already leaning towards a bistable state, but cannot evoke bistability in smaller or less bistable neurons.

**Figure 5:**
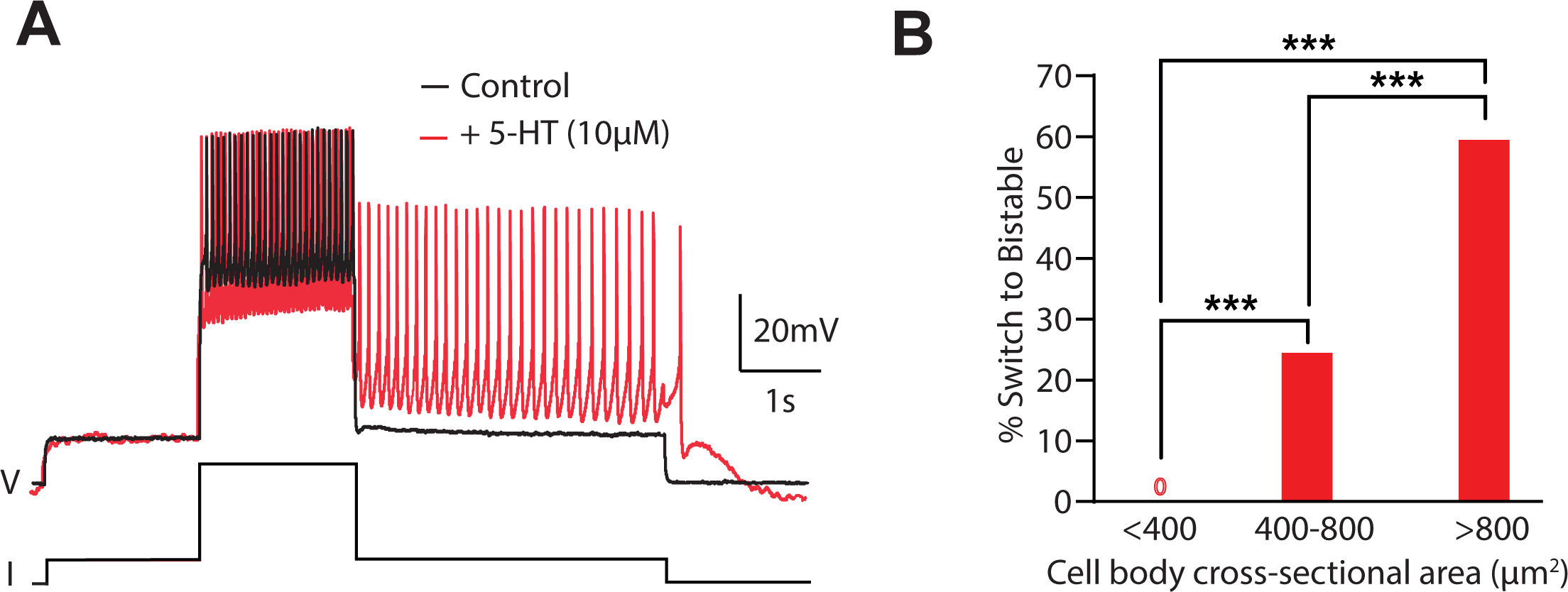
Induction of full bistability by serotonin. **A**: Non-bistable neuron’s response (top, black trace) to 2 sec depolarizing pulse (bottom). Only a small afterdepolarization is recorded after the step. During 10µM serotonin (5-HT), this neuron became fully bistable, with continuous firing (top, red trace) following the 2 sec depolarizing step (bottom). **B**: Not all non-bistable neurons respond to serotonin. Smaller neurons (<400 µm^2^) fail to show full bistability (score 4) with 5-HT. Over a quarter of intermediate-sized neurons (400-800 µm^2^) converted to full bistability during 5-HT. Most large neurons (>800µm^2^) were already bistable, and 60% of those which were not fully bistable became bistable during 5-HT. *** p<0.001 (Fisher’s exact test for **B**).

## Discussion

Our study reveals a size-based gradient in the bistable ability of α-motoneurons. Specifically, the likelihood of α-motoneurons being bistable increases with the cell body cross-sectional area. The currents, Trpm5 and INaP, which promote the active state during bistability, and Kv1, whose inactivation helps to initiate sustained firing, are more pronounced in larger α-motoneurons. Conversely, these currents are low or absent in smaller α-motoneurons. These findings highlight the importance of motoneuron size in driving bistability.

Research on bistability has primarily focused on larger α-motoneurons, identified by their ventrolateral location or by retrograde stimulation from ventral nerves (18, 31). This might have led to a failure to record the properties of smaller α-motoneurons. Neurons recorded in this study are motoneurons, as evidenced by their ventrolateral location and the expression of ChAT and Hb9 markers (Fig. 1A-B). Furthermore, they are α−motoneurons since γ-motoneurons either do not express NeuN or express it only weakly (30), and do not express GFP in Hb9-GFP mice (29, 30). Finally, virtually all of the neurons larger than 400µm^2^ also expressed MMP-9, indicative of fast α-motoneurons (32, 33). While there are no markers available to differentiate between the fast fatigue-resistant (FR) and fast fatigable (FF) α-motoneurons, we conclude that the largest neurons are FF. The smallest neurons are presumably slow (S) α-motoneurons. However, size alone is not a reliable indicator to separate these classes (42–44).

Electrophysiologically, large fast α-motoneurons differ from small slow α-motoneurons by exhibiting a delayed and accelerated firing during prolonged stimulation (31, 43, 45). Our experiments confirm this observation, as near-threshold current steps resulted in a slow depolarization in larger α-motoneurons leading to delayed firing acceleration (Fig. 2). In contrast, smaller α-motoneurons fired immediately upon reaching spike threshold followed by a spike frequency deceleration (31, 33). The depolarization and spike delay in large fast α-motoneurons were attributed to the slow inactivation of a Kv1 current (23, 33).

We now add bistability as a predominant property of large fast α-motoneurons in young mice. Over 75% of neurons over 800µm^2^ showed bistability, while only ∼10% of neurons less than 400 µm^2^ were bistable. Consistent with this, a large majority of fast α-motoneurons showed negative hysteresis during triangular ramp steps, where the derecruitment current was lower than the recruitment current, while far fewer of the slow α-motoneurons displayed negative hysteresis (33) (Fig. 2). The frequency of bistability in large α-motoneurons increased during the first weeks of postnatal development, as previously described (18); (Fig. 2C1). By P21, all large α-motoneurons were fully bistable, while smaller neurons continued to lack bistability features.

The distinct firing behaviors observed between large and small α-motoneurons can be attributed to a specific set of ionic currents that are more highly expressed in large fast α-motoneurons compared to small slow α-motoneurons. One key contributor is the activation of slow persistent inward currents (PICs), which may be carried by calcium or sodium. This activation has been linked to negative hysteresis and bistability (15, 16, 19). Expression of the PIC increases with the size of the α-motoneurons (Fig. 3B), and with the bistability score (Fig. 3D). Furthermore, our observations demonstrate the relationship between the size of the α-motoneurons and the sodium component of PICs (INaP) (Fig. 3E-H). Recently, the thermosensitive Trpm5 current has been identified as a calcium-activated sodium current driving the plateau potential in bistable mouse α-motoneurons (19). We found that the Trpm5-evoked afterdepolarization was present in virtually all large bistable α-motoneurons, but not detectable in small non-bistable α-motoneurons (Fig. 4A-C). Finally, the slowly inactivating Kv1.2-mediated current responsible for the delayed firing acceleration in large motoneurons (23) (Fig. 2), is less expressed in smaller motoneurons (Fig. 4G-I). Interestingly, this current positively correlates with the degree of bistability (Fig. 4G-I). It is likely, that the slow inactivation of Kv1 will shift the balance of currents towards depolarization, and will help to sustain continued firing in the bistable state. In sum, the lack of sufficient expression of these three pivotal currents renders small α-motoneurons non-bistable. In contrast, large α-motoneurons, which robustly express these currents, are bistable. Note that there may be some duplication of effort in these currents. Indeed, among neurons that showed continued bistable firing, 20% either did not show a marked negative hysteresis during ramps, or did not show a delayed firing acceleration. This suggest that bistability might not require the full spectrum of these currents.

Bistability in α-motoneurons was observed in early experiments only in the presence of neuromodulators (14–16, 46). However, we have revealed inherent bistability in many α-motoneurons, provided the recording temperature is sufficiently high (above 30°C) (18), to unmask the thermosensitive Trpm5 current responsible for the plateau potential (19). In our preparations, some α-motoneurons expressed intermediate properties and were not fully bistable under these conditions. Addition of 5-HT evoked full bistability in many of these neurons (Fig. 5). Interestingly, α-motoneurons completely lacking baseline bistability characteristics (bistability score 0) never became bistable with 5-HT. We propose that intermediate α-motoneurons express some of the essential currents for bistability, but at lower levels (Fig. 5). The known modulatory actions of 5-HT can then enhance these currents to sustain prolonged firing in the bistable state.

Our demonstration of a size principle for bistability contrasts with the groundbreaking work by Lee and Heckman (15, 16), who studied α-motoneurons in adult cats. In their study, they achieved a full bistability in about one-third of the α-motoneurons, which interestingly exhibited characteristics of smaller motoneurons. There are several potential explanations for the differences between our findings. First, it is uncertain whether the smallest α-motoneurons in the cat were recorded because of the use of sharp electrode recordings. Consequently, the bistability in the smallest slow α-motoneurons in cats remains unknown. Second, mouse α-motoneurons are inherently more excitable than cat α-motoneurons, primarily due to their smaller size (47, 48). Despite this, the PIC amplitude is relatively similar across both species, suggesting that PIC has a more significant impact on the firing rate in mice. Third, our observations were made on mouse α-motoneurons during the first 4 weeks of life, while Lee and Heckman’s experiments were made on adult cats. However, bistability in the larger neurons became more pronounced with age in the mouse neurons. In young adult mice (P21-P25) all large motoneurons exhibited bistability, whereas none of the smaller ones did. Finally, our results were made *ex vivo* from slices in the presence of blockers of fast synaptic transmission (though not at P21/P25) and mostly without neuromodulators. On the other hand, the cat neurons were recorded *in vivo* in the presence of methoxamine to maximize the incidence of bistability.

The role of bistability in spinal α-motoneurons remains unclear. It has been recorded during quiet standing in both rats and cats (49), suggesting a potential significance for postural control (15, 16, 26, 41). The Henneman Size Principle (50) posits that smaller α-motoneurons, due to their lower input conductance, are the first to be recruited, potentially playing a role in maintaining posture. This can be consistent with earlier work in cats, where smaller neurons showed full bistability and were possibly active during quiet standing (15, 16, 51). However, we here demonstrate, in younger mice, that small α-motoneurons are not bistable. Instead, bistability increases with the size of motoneurons (Fig. 2). Ritter et al. (51) provided evidence that large fast α-motoneurons may also be tonically recruited during quiet standing in mice. An intriguing study by Bos et al. (19) showed that mice lacking Trpm5 channels in lumbar motoneurons, exhibited compromised postural control. Given the central role of Trpm5 in bistable properties of larger α-motoneurons (19) (Fig 4), it is possible that fast α-motoneurons may play an important role in postural maintenance. The PIC plays in important role in the initiation of bistable firing, but is also critical for repetitive firing in α-motoneurons (52), and in synaptic amplification due to their dendritic location (1). Thus, the currents which together lead to bistability may individually play multiple roles in motor control in the mouse. These assumptions should be further investigated in the future using more integrated *in vivo* preparations.

## Author Contributions

R. H-W, R.B. and F.B. generated the hypotheses for the paper. R. H-W, B.D. and R.B. performed electrophysiological experiments and analysis; E.P. performed immuno-histochemical experiments and analysis. R. H-W drafted the manuscript, and R. H-W, R.B. and F.B. revised the manuscript and approved the final version of the manuscript.

## Acknowledgements

This work was supported by NIH grant NS17323 (R. H-W), by Fonds d’investissement INT FI_INT_JCJC_2019 (R.B.), by Centre National de la Recherche Scientifique (CNRS) (R.B. and F.B.) and by Agence Nationale de la Recherche Scientifique ANR-16-CE16-0004 (F.B.).

## Author notes

Correspondence: R. M. Harris-Warrick; F. Brocard; R. Bos

## References

1. Binder MD, Powers RK, and Heckman CJ. Nonlinear Input-Output Functions of Motoneurons. Physiology (Bethesda) 35: 31–39, 2020.

2. Russell DF, and Hartline DK. Bursting neural networks: a reexamination. Science 200: 453–456, 1978.

3. Russell DF, and Hartline DK. Slow active potentials and bursting motor patterns in pyloric network of the lobster, *Panulirus interruptus*. Journal of Neurophysiology 48: 914–937, 1982.

4. Hartline DK, and Graubard K. Cellular and synaptic properties in the crustacean stomatogastric nervous system. In: Dynamic biological networks: the stomatogastric nervous system, edited by Harris-Warrick RM, Marder E, Selverston AI, and Moulins M. Cambridge: M I T Press, 1992, p. 31–86.

5. Arbas EA, and Calabrese RL. Ionic conductances underlying the activity of interneurons that control heartbeat in medicinal leech. Journal of Neuroscience 7: 3945–3952, 1987.

6. Katz PS. Motor pattern modulation by serotonergic sensory cells in the stomatogastric nervous system. Cornell University, 1989.

7. Chrachri A, and Clarac F. Fictive locomotion in the fourth thoracic ganglion of the crayfish *Procambarus clarkii*. 10: 707–719, 1990.

8. Straub VA, Staras K, Kemenes G, and Benjamin PR. Endogenous and network properties of Lymnaea feeding central pattern generator interneurons. Journal of Neurophysiology 88: 1569–1583, 2002.

9. Mercer AR, Kloppenburg P, and Hildebrand JG. Plateau potentials in developing antennal-lobe neurons of the moth, Manduca sexta. Journal of Neurophysiology 93: 1949–1958, 2005.

10. Llinas R, and Sugimori M. Electrophysiological properties of in vitro Purkinje cell somata in mammalian cerebellar slices. J Physiol 305: 171–195, 1980.

11. Hultborn H, Wigstrom H, and Wangberg B. Prolonged activation of soleus motoneurones following a conditioning train in soleus Ia afferents - A case for a reverberating loop? Neurosci Lett 1: 147–152, 1975.

12. Schwindt PC, and Crill WE. Properties of a persistent inward current in normal and TEA-injected motoneurons. J Neurophysiol 43: 1700–1724, 1980.

13. Hounsgaard J, and Kiehn O. Ca^++^ dependent bistability induced by serotonin in spinal motoneurons. Experimental Brain Research 57: 422–425, 1985.

14. Hounsgaard J, and Kiehn O. Serotonin-induced bistability of turtle motoneurons caused by a nifedipine-sensitive calcium plateau potential. Journal of Physiology London 414: 265–282, 1989.

15. Lee RH, and Heckman CJ. Bistability in spinal motoneurons in vivo: systematic variations in rhythmic firing patterns. J Neurophysiol 80: 572–582, 1998.

16. Lee RH, and Heckman CJ. Bistability in spinal motoneurons in vivo: systematic variations in persistent inward currents. J Neurophysiol 80: 583–593, 1998.

17. Kiehn O, and Harris-Warrick RM. 5-HT modulation of hyperpolarization-activated inward current and calcium-dependent outward current in a crustacean motor neuron. Journal of Neurophysiology 68: 496–508, 1992.

18. Bouhadfane M, Tazerart S, Moqrich A, Vinay L, and Brocard F. Sodium-mediated plateau potentials in lumbar motoneurons of neonatal rats. J Neurosci 33: 15626–15641, 2013.

19. Bos R, Drouillas B, Bouhadfane M, Pecchi E, Trouplin V, Korogod SM, and Brocard F. Trpm5 channels encode bistability of spinal motoneurons and ensure motor control of hindlimbs in mice. Nat Commun 12: 6815, 2021.

20. Lee RH, and Heckman CJ. Essential role of a fast persistent inward current in action potential initiation and control of rhythmic firing. J Neurophysiol 85: 472–475, 2001.

21. Harvey PJ, Li X, Li Y, and Bennett DJ. 5-HT2 receptor activation facilitates a persistent sodium current and repetitive firing in spinal motoneurons of rats with and without chronic spinal cord injury. J Neurophysiol 96: 1158–1170, 2006.

22. Zhang B, Wooton JF, and Harris-Warrick RM. Calcium-dependent plateau potentials in a crab stomatogastric ganglion motoneuron. II. Calcium-activated slow inward current. Journal of Neurophysiology 74: 1938–1946, 1995.

23. Bos R, Harris-Warrick RM, Brocard C, Demianenko LE, Manuel M, Zytnicki D, Korogod SM, and Brocard F. Kv1.2 Channels Promote Nonlinear Spiking Motoneurons for Powering Up Locomotion. Cell Rep 22: 3315–3327, 2018.

24. Kiehn O, and Harris-Warrick RM. Serotonergic stretch receptors induce plateau properties in a crustacean motor neuron by a dual-conductance mechanism. Journal of Neurophysiology 68: 485–495, 1992.

25. Thuault SJ, Malleret G, Constantinople CM, Nicholls R, Chen I, Zhu J, Panteleyev A, Vronskaya S, Nolan MF, Bruno R, Siegelbaum SA, and Kandel ER. Prefrontal cortex HCN1 channels enable intrinsic persistent neural firing and executive memory function. J Neurosci 33: 13583–13599, 2013.

26. Kiehn O. Plateau potentials and active integration in the ‘final common pathway’ for motor behaviour. Trends in Neuroscience 14: 68–73, 1991.

27. Bhumbra GS, and Beato M. Recurrent excitation between motoneurones propagates across segments and is purely glutamatergic. PLoS Biol 16: e2003586, 2018.

28. Husch A, Cramer N, and Harris-Warrick RM. Long duration perforated patch recordings from spinal interneurons of adult mice. J Neurophysiol 2011.

29. Friese A, Kaltschmidt JA, Ladle DR, Sigrist M, Jessell TM, and Arber S. Gamma and alpha motor neurons distinguished by expression of transcription factor Err3. Proc Natl Acad Sci U S A 106: 13588–13593, 2009.

30. Shneider NA, Brown MN, Smith CA, Pickel J, and Alvarez FJ. Gamma motor neurons express distinct genetic markers at birth and require muscle spindle-derived GDNF for postnatal survival. Neural Dev 4: 42, 2009.

31. Manuel M, and Zytnicki D. Molecular and electrophysiological properties of mouse motoneuron and motor unit subtypes. Curr Opin Physiol 8: 23–29, 2019.

32. Kaplan A, Spiller KJ, Towne C, Kanning KC, Choe GT, Geber A, Akay T, Aebischer P, and Henderson CE. Neuronal matrix metalloproteinase-9 is a determinant of selective neurodegeneration. Neuron 81: 333–348, 2014.

33. Leroy F, Lamotte d’Incamps B, Imhoff-Manuel RD, and Zytnicki D. Early intrinsic hyperexcitability does not contribute to motoneuron degeneration in amyotrophic lateral sclerosis. eLife 3: 2014.

34. Jones HC, and Keep RF. Brain fluid calcium concentration and response to acute hypercalcaemia during development in the rat. J Physiol 402: 579–593, 1988.

35. Brocard F, Shevtsova NA, Bouhadfane M, Tazerart S, Heinemann U, Rybak IA, and Vinay L. Activity-dependent changes in extracellular Ca2+ and K+ reveal pacemakers in the spinal locomotor-related network. Neuron 77: 1047–1054, 2013.

36. Fowler SJ, and Kellogg C. Ontogeny of thermoregulatory mechanisms in the rat. J Comp Physiol Psychol 89: 738–746, 1975.

37. Harvey PJ, Li Y, Li X, and Bennett DJ. Persistent sodium currents and repetitive firing in motoneurons of the sacrocaudal spinal cord of adult rats. J Neurophysiol 2005.

38. Li X, Murray K, Harvey PJ, Ballou EW, and Bennett DJ. Serotonin facilitates a persistent calcium current in motoneurons of rats with and without chronic spinal cord injury. J Neurophysiol 97: 1236–1246, 2007.

39. Drouillas B, Brocard C, Zanella S, Bos R, and Brocard C. Persistent Nav1.6 current drives spinal locomotor functions through nonlinear dynamics. bioRxiv 2023.

40. Palmer RK, Atwal K, Bakaj I, Carlucci-Derbyshire S, Buber MT, Cerne R, Cortes RY, Devantier HR, Jorgensen V, Pawlyk A, Lee SP, Sprous DG, Zhang Z, and Bryant R. Triphenylphosphine Oxide Is a Potent and Selective Inhibitor of the Transient Receptor Potential Melastatin-5 Ion Channel. Assay Drug Dev Techn 8: 703–713, 2010.

41. Hounsgaard J, Hultborn H, Jespersen B, and Kiehn O. Bistability of 〈-motoneurons in the decrebrate cat and in the acute spinal cat after intravenous 5-hydroxytryptophan. Journal of Physiology London 405: 345–367, 1988.

42. Baczyk M, Manuel M, Roselli F, and Zytnicki D. Diversity of Mammalian Motoneurons and Motor Units. Adv Neurobiol 28: 131–150, 2022.

43. Smith CC, and Brownstone RM. Electrical Properties of Adult Mammalian Motoneurons. Adv Neurobiol 28: 191–232, 2022.

44. Thirumalai V, and Jha U. Recruitment of Motoneurons. Adv Neurobiol 28: 169–190, 2022.

45. Sharples SA, and Miles GB. Maturation of persistent and hyperpolarization-activated inward currents shapes the differential activation of motoneuron subtypes during postnatal development. eLife 10: 2021.

46. Hounsgaard J, and Kiehn O. Ca++ dependent bistability induced by serotonin in spinal motoneurons. Exp Brain Res 57: 422–425, 1985.

47. Manuel M, Chardon M, Tysseling V, and Heckman CJ. Scaling of Motor Output, From Mouse to Humans. Physiology (Bethesda) 34: 5–13, 2019.

48. Huh S, Siripuram R, Lee RH, Turkin VV, O’Neill D, Hamm TM, Heckman CJ, and Manuel M. PICs in motoneurons do not scale with the size of the animal: a possible mechanism for faster speed of muscle contraction in smaller species. J Neurophysiol 118: 93–102, 2017.

49. Eken T, Hultborn H, and Kiehn O. Possible functions of transmitter-controlled plateau potentials in alpha motoneurones. Prog Brain Res 80: 257–267; discussion 239-242, 1989.

50. Henneman E, and Mendell LM. Functional organization of motoneuron pool and its inputs. In: Handbook of Physiology The Nervous System Motor Control. Bethesda, MD: Am. Physiol. Soc., 1981, p. 423–507.

51. Ritter LK, Tresch MC, Heckman CJ, Manuel M, and Tysseling VM. Characterization of motor units in behaving adult mice shows a wide primary range. J Neurophysiol 112: 543–551, 2014.

52. Kuo JJ, Lee RH, Zhang L, and Heckman CJ. Essential role of the persistent sodium current in spike initiation during slowly rising inputs in mouse spinal neurones. J Physiol 574: 819–834, 2006.

